# The cerebellum specializes for language even in the absence of contralateral neocortical inputs

**DOI:** 10.64898/2026.08.19.745757

**Authors:** Bangjie Wang, Greta Tuckute, Hope Kean, Salomi S. Asaridou, Susan Goldin-Meadow, Susan C. Levine, Evelina Fedorenko, Anila M. D’Mello

## Abstract

Long considered a structure dedicated primarily to motor control, the cerebellum is now known to contain regions that respond selectively to language. However, how cerebellar language specialization emerges during development remains unknown. The prevailing proposal is that cerebellar functional specialization critically depends on inputs from the contralateral neocortex, received through well-established reciprocal cortico-cerebellar connections. Here, we test this proposal in two individuals with atypical brain development: EG, a right-handed woman who lacks most of her left temporal lobe (presumably from birth) and whose neocortical language network resides in her right hemisphere, and C1, an individual missing the entire left cerebral hemisphere from birth. Using precision functional MRI in EG and a cohort of 74 typically-developing adults, we find that EG’s cerebellar language network shows a strong left-hemispheric bias, mirroring the atypical lateralization of language in her cerebral cortex, while preserving canonical topography and response profiles of the language-dominant cerebellar regions. Critically, however, EG’s right cerebellar hemisphere also responds to language and even contains a language-selective region despite the absence of language regions in the neocortical left hemisphere. We replicate these cerebellar findings in C1, a more extreme case who is missing the entire left hemisphere. These results show that the cerebellum develops language sensitivity despite the absence of contralateral inputs in the face of early injury. Together, these findings challenge the view that cerebellar specialization critically requires contralateral neocortical inputs, and point instead to some degree of intrinsic neocortex-independent cerebellar organization.

## Introduction

Language is a hallmark of human cognition, which enables communication both in the moment and across generations. Historically, language has been thought to rely on the neocortex, including left temporal and frontal regions, which show substantial expansion in humans compared to other primates (e.g., Buckner and Krienen, 2013). However, the cerebellum has also expanded in the human lineage (MacLeod et al., 2003; Weaver, 2005; Balsters et al., 2010; Barton and Venditti, 2014), which likely contributed to the emergence of higher-order human skills like language. Indeed, parts of the cerebellum are robustly engaged during language tasks (Petersen et al., 1989; Stoodley and Schmahmann, 2009; LeBel and D’Mello, 2023; Turker et al., 2023), and specific subregions even show *selectivity* for language (Casto et al., 2026)—a property previously thought to be unique to the neocortex. Neuromodulation studies (Lesage et al., 2012; Argyropoulos, 2016; Turkeltaub et al., 2016; D’Mello et al., 2017; Gatti et al., 2020; Rice et al., 2021) and studies of cerebellar damage (e.g., Riva and Giorgi, 2000; Limperopoulos et al., 2014) further suggest that the cerebellum is causally important for language function.

Despite substantial recent progress in characterizing cerebellar contributions to language, *how cerebellar language regions emerge during development* remains an important open question. Cerebellar functional regions and their properties have long been argued to be inherited from the contralateral neocortex (henceforth, the *neocortical inheritance* hypothesis; Schmahmann, 1996; Strick et al., 2009). This proposal is motivated by work in non-human primates, which has established that the cerebellum is extensively anatomically connected with contralateral neocortical regions through a series of reciprocal, closed-loop circuits (Middleton and Strick, 2001; Kelly and Strick, 2003; Strick et al., 2009). These connections are thought to determine the precise locations of the different functional regions in the cerebellum and their response properties (Ramnani, 2006; Stoodley and Schmahmann, 2018).

Importantly, although the existence of regions in the human cerebellum that are functionally similar to, as well as functionally and anatomically connected with, regions in the contralateral neocortex (Buckner et al., 2011; Guell et al., 2018; Marek et al., 2018; King et al., 2023; Casto et al., 2026) is *consistent* with the idea of neocortical inheritance, it does not unambiguously support it. Further, several findings appear at odds with the neocortical inheritance hypothesis. Most importantly, the timing of when damage leads to more severe deficits sometimes differs between the cerebellum and the neocortex. For language in particular, earlier cerebellar damage is associated with more severe and longer-lasting deficits (Stoodley and Limperopoulos, 2016; Olson et al., 2023b)—the opposite of what is found in the neocortex (Müller et al., 1998; Asaridou et al., 2020; Newport et al., 2022). In addition, the relative proportions of the cerebellum dedicated to particular functions do not always mirror the proportions of the neocortex taken up by those functions (e.g., Buckner et al., 2011; Marek et al., 2018; Diedrichsen and McDougle, 2026), and some cerebellar regions exhibit functional profiles that differ from their neocortical counterparts (D’Mello et al., 2020; King et al., 2023; Shahshahani et al., 2024; Casto et al., 2026). These discrepancies raise the possibility of some degree of neocortex-independent functional organization in the cerebellum.

Here, we evaluate the neocortical inheritance hypothesis, with a focus on language, in two individuals who provide different disruptions to the expected neocortical source of cerebellar specialization—the loss of the left temporal lobe vs. the complete absence of the contralateral cerebral hemisphere. In both individuals, the neocortical language system resides in the right hemisphere (Asaridou et al., 2020; Tuckute et al., 2022; **Figure 1A**) and both have intact linguistic and cognitive abilities. By examining their cerebellar language responses, we can therefore ask whether the absence of neocortical inputs from the left hemisphere to the right cerebellum leads to the lack of specialization for language in the right cerebellum. If so, this would suggest that contralateral inputs are critically needed to set up the functions of the cerebellum (**Figure 1C, left**; typical development in **Figure 1B**). If, on the other hand, we find language regions in the right cerebellum (**Figure 1C, right**), this would tell us that such regions can arise absent contralateral neocortical inputs and thus point to some degree of intrinsic, neocortex-independent cerebellar organization. To foreshadow the key results, EG’s cerebellum—although showing left-hemispheric language dominance—also contains a language-selective region in the right hemisphere. Critically, cerebellar language responsiveness is preserved in the more extreme case, C1, who lacks her entire left cerebral hemisphere.

**Figure 1.**
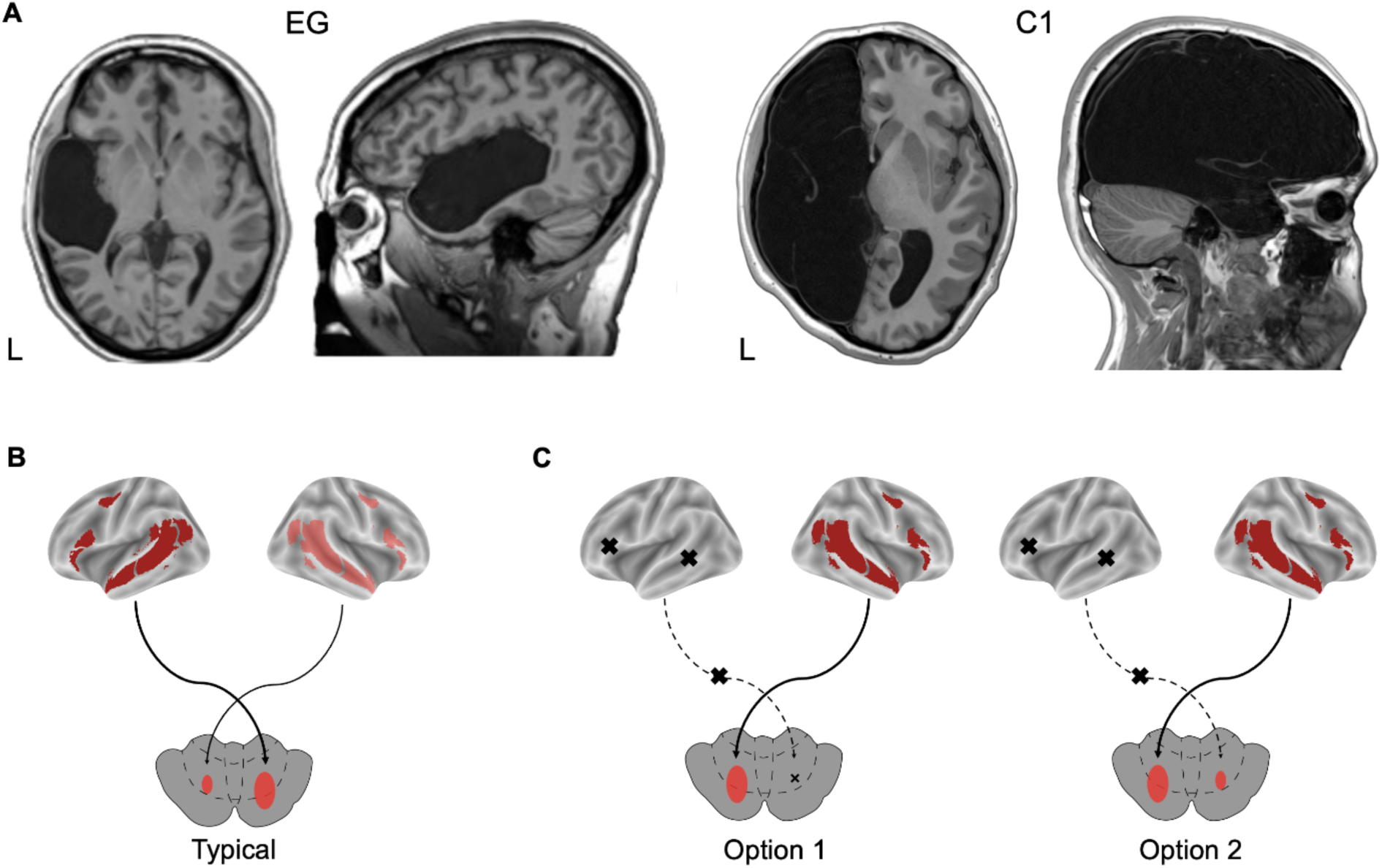
Anatomical views of EG’s and C1’s brains and an illustration of the two options for cerebellar language reorganization. **A.** T1-weighted image slices of EG’s and C1’s brains in axial and sagittal views. **B**. Typically, the neocortex shows left-lateralized language responses (top), and the cerebellum shows right-lateralized language responses (bottom). This pattern is presumed to be due to the contralateral connections between the neocortex and the cerebellum. **C.** Two possible patterns of cerebellar language reorganization in the absence of the left neocortical inputs and with the neocortical language network restricted to the right hemisphere. <u>Option 1</u>. The cerebellar language network is restricted to the left hemisphere, with no response to language in the right cerebellum. This pattern would support the neocortical inheritance hypothesis: with no input from the contralateral (left) neocortical hemisphere, no language specialization develops in the right cerebellum. <u>Option 2</u>. The cerebellar language network is left-lateralized but the right cerebellum still shows some language response and/or selectivity. This pattern would challenge the neocortical inheritance hypothesis and raise the possibility of some degree of neocortex-independent functional organization in the cerebellum.

## Results

To determine whether cerebellar language specialization depends on contralateral neocortical input, we first focused on EG, whose atypical brain organization combined with dense precision fMRI provides a uniquely informative test of this hypothesis. EG has extensive left temporal lobe loss but an intact right cerebral hemisphere that supports her language network (Tuckute et al., 2022). The precision fMRI protocol conducted in EG included a validated language task (which included a low-level control condition) and a non-linguistic task, enabling us to characterize (1) the topography of cerebellar language responses in both the left and right hemispheres, (2) whether these responses are *selective* for language, and (3) how cerebellar language response profiles compare with those of typically-developing individuals. We subsequently extend our central findings to C1, an individual missing the *entire* left cerebral hemisphere, who thus provides a more extreme test of whether cerebellar language responsiveness can persist in the absence of the expected contralateral neocortex.

### 1. Atypical, right-hemispheric neocortical language lateralization is associated with atypical, left-hemispheric cerebellar language lateralization

As EG’s neocortical language network resides in the right hemisphere (Tuckute et al., 2022; see **Supplementary Figs. 1-2** for whole-brain maps), we first asked whether this atypical neocortical language lateralization is associated with atypical cerebellar language lateralization. To do so, we used a validated language localizer paradigm based on the reading of sentences versus sequences of nonwords, which elicits robust responses at the individual-participant level (Fedorenko et al., 2010). Using activations from this paradigm, we calculated a lateralization index (LI) in EG and in each of n = 74 typically-developing (TD) control adults (see **Methods**). In contrast to the typical right-lateralized cerebellar language responses (**Figure 2A**; see **Figure 3A** for a probabilistic activation atlas from a larger number of participants—the TD group’s topography is representative of that larger sample), EG’s cerebellar language responses were strongly left-lateralized (**Figure 2B**). Considering the cerebellum as a whole, EG’s cerebellar language responses were more strongly left-lateralized than those of 73 of the 74 participants in the TD group (EG LI = 0.28 vs. TD group mean LI = -0.12, Crawford *p* = 0.008; **Figure 2C**). EG also showed some response to language in her right cerebellum, as we discuss in detail in the penultimate section of the Results.

**Figure 2.**
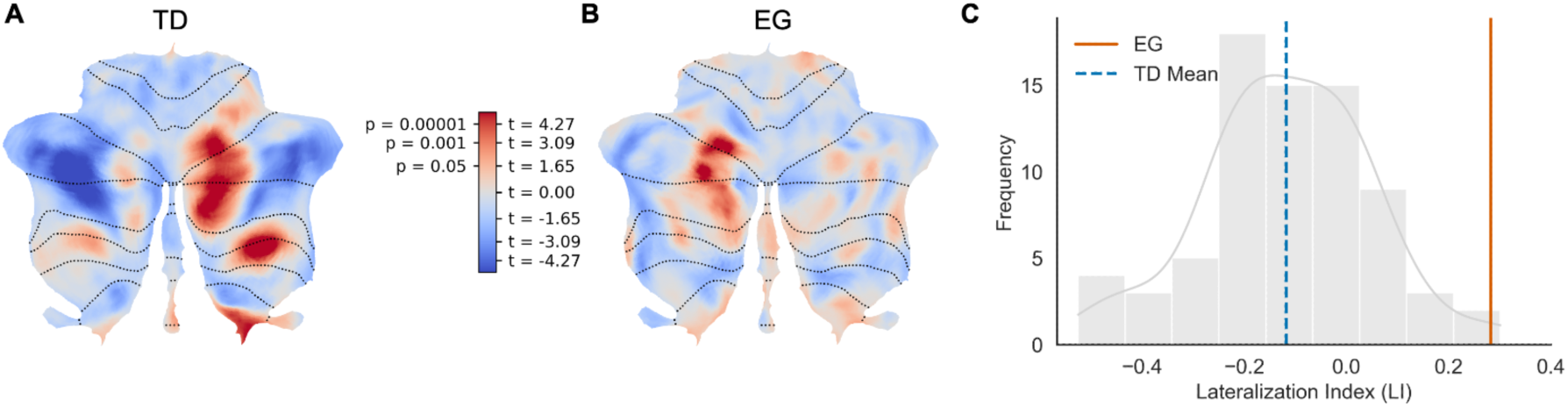
Lateralization of cerebellar language responses in an individual with an atypically lateralized (likely due to the loss of the left temporal lobe) neocortical language network. Cerebellar responses to language (Sentences > Nonwords) in n = 74 TD participants (**A**) and in EG (**B**) show that language is right-lateralized in the cerebellum of TDs and left-lateralized in EG. These hemispheric biases are evident in both the activation maps (A and B) and in the distribution of the lateralization indices (LIs) in TD language activations (gray bars; **C**), with the mean TD LI shown in blue, and EG’s LI shown in orange.

**Figure 3.**
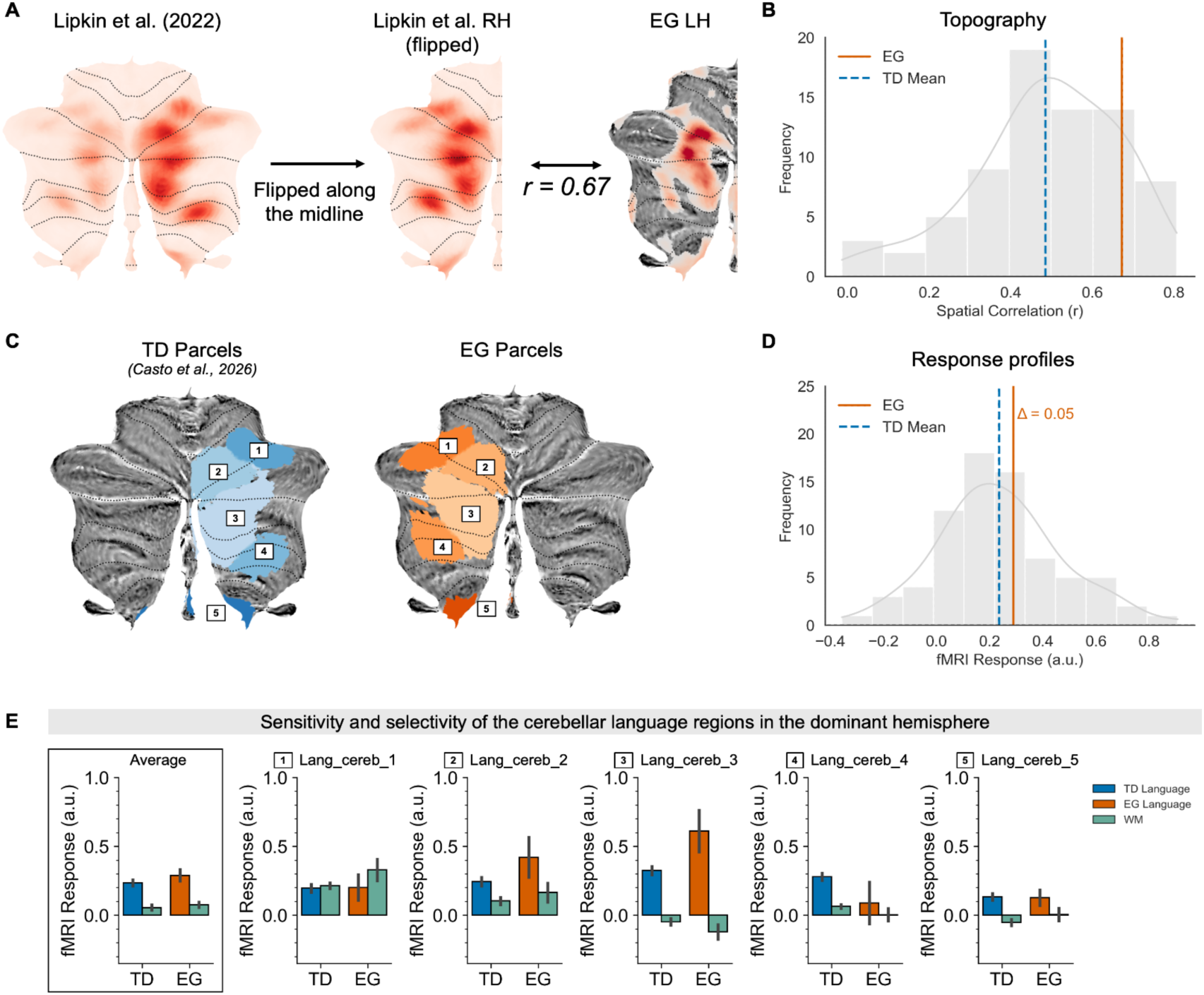
The topography and response profiles of language-dominant cerebellar regions in EG vs. TDs. **A**. Voxel-wise spatial correlation between EG’s language-dominant cerebellar language activation pattern and the RH cerebellar probabilistic activation atlas created from 806 TD adults (Lipkin et al., 2022). Note that the TD probabilistic atlas has been flipped along the midline to calculate voxel-wise spatial correlations. **B**. Distribution of spatial correlations between each TD participant’s language-dominant activation pattern and the RH cerebellar probabilistic activation atlas (gray bars). The spatial correlation for the TD group mean is shown in blue and EG’s spatial correlation is shown in orange. **C.** Cerebellar parcels for TDs and EG in the language-dominant cerebellar hemisphere (EG’s parcels are the same as TDs, just flipped onto the left hemisphere). The parcels and their numbering come from Casto et al. (2026). **D**. Distribution of response magnitudes for the language contrast (Sentences > Nonwords; averaged across the five cerebellar regions) in TD participants (gray bars), with the mean TD response magnitude shown in blue and EG’s language-dominant (LH) response magnitude shown in orange. **E**. Responses to the language (Sentences > Nonwords) and spatial WM (Hard > Easy) contrasts in the language-dominant cerebellar hemisphere (LH for EG, RH for TD individuals) in each of the five regions and averaged across them (box). EG’s language responses are shown in orange, TD participants’ language responses are shown in blue, and the WM responses for both EG and TDs are shown in green. For EG, the error bars indicate the standard error of the mean by experimental runs; for the TD group, the error bars indicate the standard error of the mean by participants.

### 2. EG’s atypically lateralized language-dominant (left) cerebellum preserves a typical topography and response profile

We next asked whether EG’s language responses in the left cerebellum retained (a) typical topography, and (b) typical response profiles of the language-dominant cerebellum despite the shift in lateralization. To assess the typicality of EG’s cerebellar language topography, we computed voxel-wise spatial correlations between each participant’s language-dominant cerebellar activation map (left cerebellum in EG, right cerebellum in TDs) and the right cerebellum in a probabilistic activation atlas derived from n = 806 TD participants (Lipkin et al., 2022; **Figure 3A**). The topography of the language responses in EG’s language-dominant cerebellum showed strong correspondence with the typical language topography (*r* = 0.67; **Figure 3A**). This correlation value did not significantly differ from the TD distribution (TD group mean = 0.49, Crawford *p* = 0.15; **Figure 3B**).

To examine whether the response profiles of EG’s language-dominant cerebellar language regions remained typical, we next estimated response magnitudes to each condition of the language localizer (Sentences, Nonwords) and to each condition of a non-linguistic demanding spatial working memory task (Hard, Easy; see **Methods**) in individually-defined language functional regions of interest (fROIs). To define subject-specific cerebellar language fROIs, we used five parcels, derived from a probabilistic activation atlas, corresponding to distinct regions within the language-dominant (right) cerebellar hemisphere (Casto et al., 2026; **Figure 3C**). For EG, these parcels were flipped onto the left cerebellar hemisphere. For each participant, within each parcel, we used a portion of the language localizer data to identify the most language-responsive voxels, and then estimated the responses to the language and spatial working memory conditions in held-out data (see **Methods**). For each fROI, the magnitude of the language contrast (Sentences > Nonwords) can be used as an index of language *sensitivity*, and the magnitude of the non-language contrast (Hard > Easy spatial working memory) can be used to characterize the *selectivity* for language relative to non-linguistic tasks (see Casto et al., 2026 and Wolna et al., 2026 for a similar approach).

As previously reported, in TD individuals, all language regions in the language-dominant cerebellum show strong language *sensitivity*, and a subset further shows a stronger response to language compared to non-linguistic stimuli and tasks (language *selectivity*) (Fedorenko et al., 2010; Lipkin et al., 2022; Casto et al., 2026; Wolna et al., 2026). Averaging across the five language regions, the magnitude of the Sentences > Nonwords contrast was similar between EG and TDs (EG language response = 0.29 vs. TD group mean = 0.24; Crawford *p* = 0.41; **Figure 3D,E**).

Overall, EG’s pattern of language selectivity was also similar to that of TDs (**Figure 3E**). In particular, Lang_cereb_3 was the most language-selective region in both EG and TDs, with a strong response to language and no response to the working memory task. Lang_cereb_5 also showed a selective profile in both EG and TDs, with no response to the working memory task, although the response to language was relatively low. Two of the remaining regions— Lang_cereb_1 and Lang_cereb_2—showed a mixed-selective response in both EG and TDs, with a relatively strong response to the spatial working memory task in addition to language. The last region—Lang_cereb_4—showed a relatively selective response in TDs, but no clear response to either language or spatial working memory in EG.

### 3. Despite the lack of left-hemispheric neocortical inputs, EG’s right cerebellum responds to language and contains a language-selective region

Critically, as foreshadowed above, in addition to the largely typical-like responses to language in her language-dominant (left) cerebellum, EG also showed some response to language in her right cerebellum. Similar to the language-dominant hemisphere, we evaluated the typicality of the topography and response profiles of EG’s cerebellar language regions, except here, we compared EG’s right cerebellum to the TD left cerebellum. The topography only minimally resembled the TD topography (*r* = 0.28; **Figure 4A**). Although this value was lower than that of ∼65% of the TD population, it did not differ significantly from the TD distribution (TD group mean = 0.35, Crawford *p* = 0.35; **Figure 4B**). Note also that, in general, the topography of language responses in the language non-dominant cerebellar hemisphere is more variable across individuals than in the language-dominant hemisphere.

**Figure 4.**
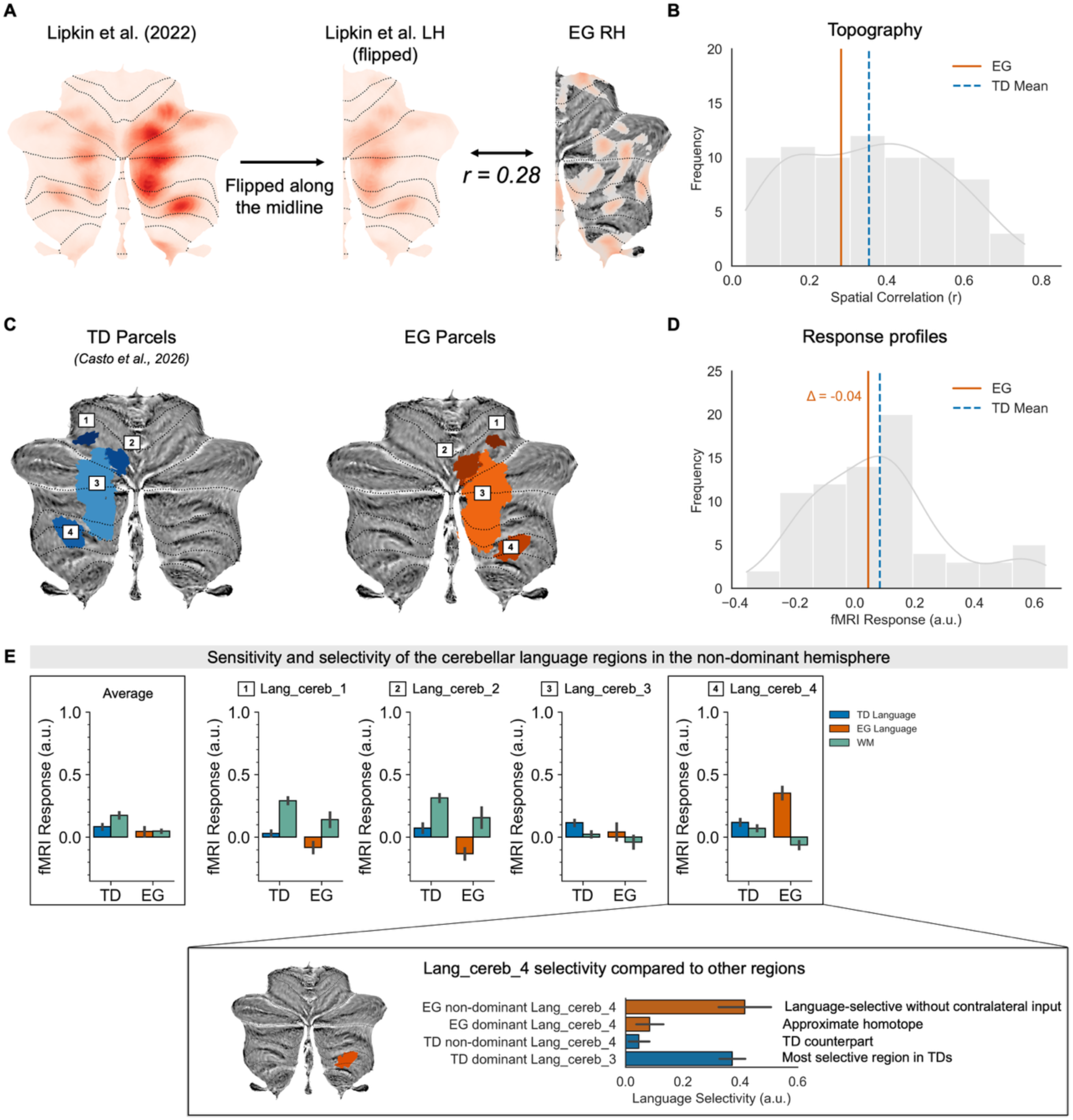
The topography and response profiles of language non-dominant cerebellar regions in EG vs. TDs. **A**. Voxel-wise spatial correlation between EG’s language non-dominant cerebellar language activation pattern and the LH probabilistic activation atlas created from 806 TD adults (Lipkin et al., 2022). Note that the TD probabilistic atlas has been flipped along the midline to calculate voxel-wise spatial correlations. **B**. Distribution of spatial correlations between each TD participant’s language non-dominant activation pattern and the LH cerebellar probabilistic activation atlas (gray bars). The spatial correlation for the TD group mean is shown in blue and EG’s spatial correlation is shown in orange. **C.** Cerebellar parcels for TDs and EG in the language non-dominant cerebellar hemisphere (EG’s parcels are the same as TDs, just flipped onto the right hemisphere). The parcels and their numbering come from Casto et al. (2026). **D**. Distribution of response magnitudes for the language contrast (Sentences > Nonwords; averaged across the four cerebellar regions) in TD participants (gray bars), with the mean TD response magnitude shown in blue and EG’s language non-dominant (RH) response magnitude shown in orange. **E**. Responses to the language (Sentences > Nonwords) and spatial WM (Hard > Easy) contrasts in the language non-dominant cerebellar hemisphere (RH for EG, LH for TD individuals) in each of the four regions and averaged across them (box). EG’s language responses are shown in orange, TD participants’ language responses are shown in blue, and the WM responses for both EG and TDs are shown in green. For EG, the error bars indicate the standard error of the mean by experimental runs; for the TD group, the error bars indicate the standard error of the mean by participants. **Inset**. Language selectivity in EG’s language non-dominant (right) Lang_cereb_4 compared to other parcels including EG’s language-dominant Lang_cereb_4, TDs’ language non-dominant Lang_cereb_4, and TDs’ language-dominant Lang_cereb_3. Language selectivity is computed as the fMRI response to the Sentences > Nonwords contrast in the language task minus the fMRI response to the Hard > Easy contrast in the spatial WM task.

With respect to the response profiles, in TD individuals the language regions in the language non-dominant cerebellum (**Figure 4C**; note that these regions are not homotopes of language-dominant cerebellar regions) show language *sensitivity*, but only Lang_cereb_3 shows some degree of language *selectivity* (**Figure 4E**, in line with Casto et al., 2026). Averaging across the four language regions, the magnitude of the Sentences > Nonwords contrast was low and similar between EG and TDs (EG language response = 0.04 vs. TD group mean = 0.08; Crawford *p* = 0.43; **Figure 4D,E**).

The pattern of language selectivity was similar between EG and TDs for Lang_cereb_1 and Lang_cereb_2: in both EG and TDs, these regions were non-selective, showing stronger responses to the working memory contrast than to language (**Figure 4E**). For Lang_cereb_3, TDs but not EG showed a reliable response to language, although EG’s response magnitude to language in this region did not reliably differ from the TD distribution (Crawford *p* = 0.39). Note that this lack of a reliable language response in EG is in contrast to the language-dominant cerebellum, where the approximately homotopic Lang_cereb_3 showed the strongest and most selective response to language in EG, similar to TDs. Finally, and most excitingly, Lang_cereb_4 showed a strongly selective language response in EG, in contrast to a mixed-selective response in TDs. The selectivity of this region in EG was higher than that of approximately 80% of TDs for the same region (though EG’s language selectivity was still within the TD distribution, Crawford *p* = 0.19), and was reliably stronger than that in EG’s approximately homotopic Lang_cereb_4 region in the language-dominant hemisphere (**Figure 4E inset**).

Two other aspects of the data are worth noting. First, the selectivity of EG’s Lang_cereb_4 was comparable to the selectivity in TDs’ most language-selective region (Lang_cereb_3) in the language-dominant cerebellar hemisphere (Crawford *p* = 0.45; **Figure 4E inset**). And second, the fact that EG’s left Lang_cereb_4 (the approximate homotope in the language-dominant cerebellar hemisphere) was not language-selective (EG language selectivity = 0.09; cf. the selectivity of her non-dominant Lang_cereb_4 = 0.42) makes it unlikely that Lang_cereb_4 in her language non-dominant hemisphere inherits its language selectivity from the left cerebellum.

### 4. The right cerebellum responds to language even in an individual missing the entire left hemisphere

A caveat regarding the current evaluation of the neocortical inheritance hypothesis in EG is that she still retains tissue in the left neocortex. While this tissue was not language-responsive (**Supplementary Fig. 2**; Tuckute et al., 2022), it is nonetheless possible that remaining left neocortical regions send lower-level linguistic inputs (e.g., acoustic and phonemic information) to support the functional specialization of the right cerebellum. To address this, we examined whether cerebellar functional specialization can emerge in the complete absence of the contralateral neocortex. At the time of scanning, C1 was a 14-year-old female missing the entire left hemisphere from birth, with no cognitive or linguistic impairment, who lateralizes language completely to the right neocortical hemisphere (**Supplementary Fig. 3A**; also see **Methods**; Asaridou et al., 2020). C1’s cerebellum showed a leftward lateralization for spoken stories (**Figure 5A-5B**, note that the contrast used to examine language responses in C1 was Stories > baseline, which differed from the Sentences > Nonwords contrast used in EG; also see **Supplementary Fig. 3B** for additional lateralization results). More importantly, despite no contralateral inputs, the topography of language responses in C1’s right cerebellum showed strong correspondence with the typical language topography in the right cerebellum (*r* = 0.72; **Figure 5B inset**). Further, several regions in C1’s right cerebellum showed robust responses to language (here, spoken stories) (**Figure 5C**; see **Supplementary Fig. 3C** for LH cerebellar results in C1), including high responses to language in Lang_cereb_3 (TDs’ most language-responsive and selective cluster).

**Figure 5.**
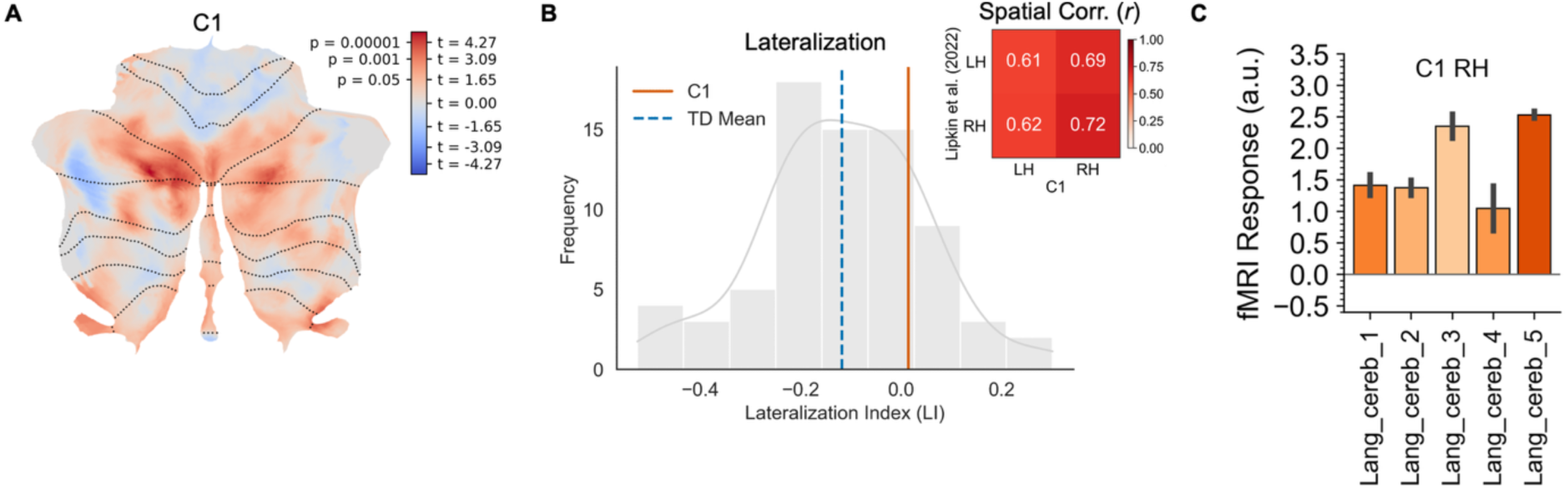
The topography and response profiles of the right cerebellar hemisphere in C1. **A.** Cerebellar responses to language (Spoken Stories > baseline) in C1. **B.** Distribution of the lateralization indices (LIs) in TD language (Sentences > Nonwords) activations (gray bars), with the mean TD LI shown in blue and C1’s language (Spoken Stories > baseline) LI shown in orange. **Inset.** Voxel-wise spatial correlations between C1’s cerebellar activation patterns and the cerebellar probabilistic activation atlas in Lipkin et al. (2022). **C.** Responses to the language (Spoken Stories > baseline) contrast in C1’s right cerebellar regions. Given that C1’s RH language activation topography more closely resembled a typical RH, the responses were extracted from TDs’ language-dominant (right) cerebellar parcels. The error bars indicate the standard error of the mean by experimental runs.

## Discussion

How the cerebellum develops specialization for language remains debated and—short of longitudinal early-childhood studies—is challenging to understand in typical brains. To evaluate proposals about the emergence of language-specialized cerebellar regions, we turned to two individuals with atypical brains: EG is missing most of her left temporal lobe, likely from birth, and shows right-lateralized neocortical responses to language, in the presence of intact language and cognitive functioning (Tuckute et al., 2022). The lack of neocortical language responses in EG’s left hemisphere allowed us to evaluate the claim that contralateral neocortical inputs critically guide cerebellar functional organization (Schmahmann, 1996; Strick et al., 2009). We observed that (1) EG’s cerebellum showed left-hemispheric language dominance, mirroring the right-hemispheric dominance of her neocortical language network, and (2) EG’s language-dominant cerebellar regions were topographically and functionally similar to typical language-dominant cerebellar regions. Critically, we also found that (3) EG’s language non-dominant right cerebellar hemisphere responded to language and even contained a language-selective region. We replicate the finding of right cerebellar language responsiveness in the absence of contralateral inputs in a second individual, C1, who is missing the entire left neocortical hemisphere from birth. Together, these findings provide evidence that cerebellar specialization for language can arise without input from the contralateral neocortical language regions. Below, we first consider what EG reveals about the organization of cerebellar language representations, leveraging the precision of her imaging to characterize this organization in detail, and then turn to C1 to ask whether the central finding extends to the more extreme case of complete absence of the contralateral cerebral hemisphere.

EG’s cerebellar language system showed atypical leftward lateralization for language, mirroring the atypical rightward lateralization of her neocortical language network, and looked like a typical language-dominant cerebellum. The overall topography was conserved, and the response profiles of EG’s left cerebellar language regions were similar to those of the corresponding regions in typically-developing (TD) participants’ right cerebellum. These results suggest that the hemispheric bias for language can flip in the cerebellum, much like what is often observed in the neocortex after early damage to the left hemisphere (Asaridou et al., 2020; Tuckute et al., 2022). In a different task paradigm (naturalistic listening), C1’s left cerebellum also showed a relatively typical pattern of language responses—closer topographically to TDs’ dominant cerebellar hemisphere than to TDs’ non-dominant cerebellar hemisphere. The fact that language function is intact in both cases further suggests that the atypically lateralized left cerebellar language regions can support language function, although we do not have evidence of their causal contributions. Much past work has emphasized the early equipotentiality for language of the two neocortical hemispheres (Asaridou et al., 2020; Tuckute et al., 2022; Martin et al., 2024). To our knowledge, a similar claim has not been made for the cerebellum, and our data point in this direction.

Future work should aim to understand the determinants of the cerebellum’s language dominance. For example, does it always ‘track’ neocortical language dominance? Some past studies (Jansen et al., 2005; Berl et al., 2014; Casto et al., 2026) have reported a correlation between the degree of neocortical vs. cerebellar language lateralization, but variance in the degree of cerebellar language dominance is not fully explained by the neocortical bias. For instance, despite right-lateralized neocortical language responses and lack of the left neocortex, C1’s cerebellar language responses were relatively bilateral (although more language-responsive voxels were still found on the left compared to the TD group). A more complete understanding of this relationship, including tracking changes therein across the lifespan and in atypical brains, can help to illuminate the influence of the neocortex on lateralization in the cerebellum or vice versa.

Another more general question for future work is whether/to what extent the neocortex (and which particular neocortical regions) is necessary for the emergence of typical language organization and responses in the cerebellum, and intact language function. Our cases span a continuum of left-hemisphere loss, from more focal absence of the left temporal lobe to complete absence of the left cerebral hemisphere, with both individuals showing language-responsive regions in the cerebellum. While EG’s right cerebellum was topographically dissimilar from that of TDs, C1’s right cerebellum was more topographically similar. An interesting direction for future research is to establish whether the extent of early neocortical loss influences the degree to which cerebellar organization is preserved, with more extensive loss potentially permitting more canonical organization. One possibility is that the residual left neocortical tissue in EG provided atypical, non-linguistic input to the right cerebellum, which could be more disruptive to the emergence of typical functional organization than no input at all. Some additional insights may come from individuals with substantially reduced or malformed neocortex and white matter (for example, due to severe hydrocephalus; Lewin, 1980; Feuillet et al., 2007; Forsdyke, 2015; cf. Neuroskeptic, 2015). We are not aware of anyone having examined the cerebellum’s functional organization and its contributions to language and cognition in this population.

Critically, despite the absence of language inputs from the contralateral neocortex, the right cerebellum in both EG and C1 shows responses to language and, in EG, even some degree of language selectivity in one of the regions. These results demonstrate that inputs from functionally-specialized contralateral neocortical regions are not required to develop functional specialization in the cerebellum, challenging the widely-held view that the neocortex guides cerebellar organization (Schmahmann, 1996; Strick et al., 2009)—what we referred to as the neocortical inheritance hypothesis. It is worth noting that our findings align with prior work: even though cerebellar language responses are typically not discussed, prior fMRI investigations of individuals with anatomically atypical left neocortical hemispheres and right-hemispheric neocortical language dominance also show language responses in both cerebellar hemispheres, including in individuals missing varying degrees of the left hemisphere (Asaridou et al., 2020; Newport et al., 2022).

How do responses to language and language specialization arise in the right cerebellum in the absence of contralateral neocortical language inputs? Although structurally intact, EG’s frontal lobe did not show any response to language (Tuckute et al., 2022). We did observe some activation for the language localizer contrast in EG’s left angular gyrus; however, the responses to the language condition were not above the low-level fixation baseline, and these clusters showed low/no functional connectivity with the right-hemisphere language regions (see **Supplementary Figs. 1-2**; **Supplementary Method Section 1**). It is therefore unlikely that the remaining tissue in EG’s left hemisphere contributed to the strong and selective responses to language in the right cerebellum. Furthermore, C1 provides a more definitive example: unlike EG, C1 lacks the entire left cerebral hemisphere, eliminating not only residual left-hemisphere language cortex but also any left neocortical substrate through which contralateral neocortical inputs could influence cerebellar organization. Despite this, C1’s right cerebellum remained responsive to language.

We consider three alternative possibilities for how language responses (and even selectivity in the case of EG) may arise absent inputs from the contralateral neocortical hemisphere: ipsilateral neocortical inputs, inputs from the language-dominant cerebellar hemisphere, and intrinsic organization of the cerebellum.

Could the right cerebellum receive some language inputs from the *ipsilateral* neocortex? Although cerebro-cerebellar pathways are overwhelmingly crossed (estimates ranging from 60 to 80% depending on species and target regions; Schmahmann, 1996; Serapide et al., 2002; Henschke and Pakan, 2020), and show this dominant contralateral pattern already in utero (Pieterman et al., 2017), tract-tracing studies in monkeys and rats do provide some evidence of ipsilateral cerebro-cerebellar connections (Wiesendanger and Wiesendanger, 1985; Schmahmann and Pandya, 1989; Cicirata et al., 2005; Na et al., 2019). Unfortunately, these studies rarely characterize the full circuit (neocortex to the cerebellum), often examining thalamo-cerebellar or ponto-cerebellar circuits as proxies (Rosina and Provini, 1984), and it is challenging to extrapolate from these animal models to humans given substantial differences in cerebellar and neocortical anatomy and cytoarchitecture (Busch and Hansel, 2023). Nevertheless, in principle, some ipsilateral connections may exist in humans between the language regions in the neocortex and some parts of the cerebellum. The main challenge for this possibility in EG’s data is that the organization of the right cerebellar language responses in EG differs substantially from the organization of her language-dominant (left) cerebellum. If inputs from the same (right) neocortical hemisphere set up *both* of the cerebellar hemispheres, one would expect a more symmetrical cerebellar language system. Instead, the overall topography and the location of the most language-selective region vary between EG’s two cerebellar hemispheres (it is Lang_cereb_3 in the left cerebellum vs. Lang_cereb_4 in the right cerebellum). Similarly, while C1’s cerebellar language responses were more bilaterally distributed than in EG, they were nonetheless not exactly symmetrical.

A second possibility is that the right cerebellum develops language responses and selectivity via inputs from the left cerebellum. In sharp contrast to the neocortex, the cerebellum possesses very few *direct* connections between its hemispheres, with no compelling evidence to date of direct anatomical connections between the left and right dentate nuclei (the output nuclei of the cerebellar cortex) (Mihailoff, 1983; Eccles, 2013; Houck and Person, 2014). Some communication may occur through closed-loop interactions with extracerebellar structures, such as the pons (Rosina and Provini, 1984; although see above for relative sparsity of ipsilateral connections that would then be required). However, as with the first possibility introduced earlier, the fact that the organization of the right cerebellum does not mirror that of the left cerebellum casts doubt on this possibility.

Finally, and perhaps most likely, the right cerebellum may exhibit some intrinsic responses to language and some degree of self-organization. In fact, early researchers of the cerebellum were quite open to this possibility. For example, in their foundational work, Snider and Eldred (1952) discuss the existence of sensory representations in both the neocortex and the cerebellum (e.g., “with recognition of tactile, auditory, and visual regions in both the cerebrum and cerebellum it is logical to question whether or not these regions are interrelated, and if so, what the functional significance is of such an interrelationship”). Against this backdrop, the discovery of cerebro-cerebellar connections was almost surprising: Snider and Eldred note that such connections turned out to be “more extensive … than hitherto considered possible.” And, as noted in the Introduction, the existence of such connections is *compatible* with the neocortex driving the cerebellum’s organization but does not unambiguously support the direction of this relationship: such connections are also consistent with the cerebellum driving neocortical organization, or with independent self-organization in the neocortex and the cerebellum. Distinguishing among these possibilities critically requires longitudinal functional imaging data, including data from early development.

If we are to take seriously the possibility that language function emerges in the cerebellum without neocortical inputs, what other inputs could drive this emergence? Studies in cats have shown that the cerebellum receives inputs from the superior olivary complex, the inferior colliculus, and the cochlea, with auditory response latencies similar to those in primary auditory cortex; these responses are present even in decerebrate animals (Snider and Stowell, 1944; Altman et al., 1976; Huang et al., 1982). These direct auditory inputs may set up primary auditory representations in the cerebellum, which are present in humans (Petacchi et al., 2005), and could provide access to the auditory information that likely scaffolds the development of the language regions in hearing individuals. Consistent with this intriguing possibility, neuroimaging studies suggest that the cerebellum is functionally organized remarkably early, with many large-scale networks detectable in newborns and showing relative stability across the early years of life. Notably, these early cerebellar networks emerge before long-range cerebello-neocortical connections are fully established (Kipping et al., 2017; Lyu et al., 2025; Tikoo et al., 2025).

Although intriguing, the results of our study should be considered in light of the study’s limited scope. First, this is a study with two cases, and extending the findings to other individuals remains an important goal. Second, the cases differed in age and task paradigms, providing complementary but not directly comparable evidence. Third, our study—like most studies in the field—examines the brain at a particular time point (adulthood in EG, adolescence in C1), and making inferences about brain development from a mature brain’s architecture is not straightforward and should ideally be supplemented by longitudinal studies matched on tasks and demographics. Finally, we have focused here on the language circuits; whether the findings generalize to other cognitive functions also remains to be determined.

In summary, our findings of language responses and language selectivity in the cerebellar hemisphere that lacks or has disrupted contralateral neocortical language inputs (1) pose a challenge to the prominent view whereby neocortical inputs set up cerebellar organization, (2) suggest some degree of neocortex-independent cerebellar organization, possibly driven by direct sensory inputs from conserved structures in the midbrain and brainstem, and (3) highlight just how little is currently known about the neural architecture required to support speech and language in the human brain. A key theoretical implication is that accounts of the emergence of speech and language in the infant brain should become less cortico-centric given the intriguing possibility of parallel, neocortex-independent emergence of the relevant circuits in the cerebellum. A practical corollary is that the current wave of developmental neuroscience work, including in newborns and infants (Ellis and Turk-Browne, 2018; Kosakowski et al., 2024; Olson et al., 2025; Behm et al., 2026; O’Doherty et al., 2026), should ensure that the cerebellum is not “cut off” in the field of view of MRI acquisition sequences (a common problem in much prior and ongoing neuroimaging research; Wang et al., 2025): even if the current studies are focusing on neocortical responses, inclusion of the cerebellum will afford exciting possibilities for later re-analysis of these incredibly precious and challenging-to-collect datasets. Finally, our study highlights the importance of anatomically atypical brains, even in the context of single-case studies, in furthering our understanding of general constraints on the anatomy and functional organization of the brain.

## Methods

### Participants

#### Participants of interest

The current study focused on two participants of interest, EG and C1. Detailed information on EG has been previously reported by Tuckute et al. (2022). Briefly, EG is a right-handed female with no motor, language, or other cognitive deficits. EG’s left temporal lobe has likely been missing from birth (**Figure 1A**). The etiology of the anatomical anomaly is difficult to establish unambiguously, but the most likely scenario is a congenital cyst, perhaps following a prenatal/perinatal stroke. She did not have any head trauma or injuries as a child or adult. A detailed examination of the cerebellum carried out for this study additionally revealed a small cyst in EG’s cerebellar vermis. This finding appeared benign and was unlikely to affect the reported results or interpretation. EG took part in several fMRI sessions at MIT across a series of visits, two of which (Session 1, 2016, 54 years; Session 2, 2021, 59 years) are analyzed in the present study.

Detailed information on C1 has been previously reported by Asaridou et al. (2020). Briefly, C1 is a left-handed female, possibly as a consequence of right hemiparesis. Apart from the hemiparesis, she shows no language or other cognitive deficits. C1’s left neocortical hemisphere was absent from birth due to prenatal left internal carotid occlusion. C1 was recruited from a major city and scanned at an academic hospital when she was 14 years old. Her parents gave written informed consent following the guidelines of the Institutional Review Boards of the Division of Biological Sciences at The University of Chicago, and the Office of Research at the University of California, Irvine, which approved the study. C1 gave verbal assent.

#### Neuroanatomically typical control participants

Data from 74 typically-developing (TD) individuals (age range = 18-60 years, mean age = 26, standard deviation = 6.71; 39 females) were analyzed. Seventy participants were right-handed; the remaining four participants were left-handed or ambidextrous. All participants were native speakers of English or highly proficient in English and had normal hearing and normal (or corrected-to-normal) vision.

EG and TD participants were recruited from the greater Boston area, scanned at the Massachusetts Institute of Technology (MIT), and compensated for their time. They gave informed written consent in accordance with the requirements of MIT’s Committee On the Use of Humans as Experimental Subjects.

### Functional MRI tasks

All participants scanned at MIT completed several functional MRI tasks. Here, we describe two tasks that were analyzed for the current project: a language localizer task (Fedorenko et al., 2010), and a spatial working memory task, which is commonly used as a localizer for the multiple demand, or executive control, network (Blank et al., 2014; Assem et al., 2020a; Shashidhara et al., 2020), and which was used here to evaluate the selectivity of the language regions for language vs. general cognitive demand.

#### Language localizer task

This localizer was introduced in Fedorenko et al. (2010) and has been used in many subsequent studies (e.g., Blank et al., 2016; Fedorenko et al., 2020; Chen et al., 2023; Hu et al., 2023; Shain et al., 2024; Tuckute et al., 2024; the task is available for download from https://www.evlab.mit.edu/resources). Participants silently read sentences and lists of unconnected, pronounceable nonwords in a blocked design. The Sentences > Nonwords contrast targets cognitive processes related to high-level language comprehension, including understanding word meanings and combinatorial linguistic processing. This contrast robustly activates the neocortical fronto-temporal network and its right-hemisphere homotopic regions, as well as additional neocortical and subcortical regions (Wolna et al., 2026), including, critically, several regions in the cerebellum (Casto et al., 2026). Importantly, the Sentences > Nonwords contrast generalizes across presentation modalities (e.g., reading vs. listening vs. audiovisual processing), types of tasks, stimuli within a language (e.g., sentences vs. passages, naturalistic vs. hand-constructed), and diverse languages (e.g., Fedorenko et al., 2010; Scott et al., 2017; Malik-Moraleda et al., 2022; Chen et al., 2023; Olson et al., 2023a; Gao et al., 2026; see Fedorenko et al., 2024a, 2024b for a review). In addition, a system that closely corresponds to the one activated by the language localizer can be found in patterns of functional connectivity (Braga et al., 2020; Du et al., 2024, 2025; Shain and Fedorenko, 2025), although this approach typically requires more data for reliable within-individual identification compared to the localizer.

Each stimulus (Sentence or Nonword list) was 6 s long and consisted of 12 words or nonwords presented one word/nonword at a time at the rate of 450 ms per word/nonword. The main task was attentive reading. Each stimulus was followed by a simple button-press task, which was included to maintain alertness. Trials were grouped into blocks of 3 trials of the same condition (18 s total). Each scanning run consisted of 16 blocks (8 per condition) and 5 blocks of a baseline condition (blank screen) (14 s each), for a total run duration of 358 s. Condition order was counterbalanced across runs. Each participant completed 2 runs.

#### Spatial working memory (WM) task

The spatial WM task was introduced in Fedorenko et al. (2011) and has been used in many subsequent studies as a localizer for the multiple demand system (Blank et al., 2014; Shashidhara et al., 2019, 2020, 2024; Diachek et al., 2020; Malik-Moraleda et al., 2022). Participants had to keep track of spatial locations presented in a sequence within a grid (8 locations in the Hard condition, 4 locations in the Easy condition) in a blocked design. The Hard > Easy contrast targets cognitive processes broadly related to performing demanding tasks—what is often referred to by the umbrella term ‘executive function’. Importantly, the Hard > Easy spatial WM contrast generalizes to other contrasts of more vs. less demanding conditions (e.g., Duncan and Owen, 2000; Fedorenko et al., 2013; Hugdahl et al., 2015; Shashidhara et al., 2019; Assem et al., 2020b), and a system that closely corresponds to the one activated by the multiple demand localizer can be found in patterns of functional connectivity (e.g., Buckner et al., 2011; Assem et al., 2020b; Braga et al., 2020; Du et al., 2024). This and similar contrasts robustly activate a bilateral neocortical fronto-parietal network, and several regions in the cerebellum (Marek et al., 2018; Braga et al., 2020; Casto et al., 2026).

Each trial consisted of a brief fixation cross shown for 500 ms followed by 4 sequential flashes of unique locations within a 3 × 4 grid (1 s per flash; two locations at a time in the Hard condition, one location at a time in the Easy condition). Each trial ended with a two-alternative forced-choice question (two sets of locations were presented for up to 3.25 s, and participants had to choose the set of locations they just saw; if they responded before 3.25 s elapsed, there was a blank screen for the remainder of the 3.25 s period). Participants were given feedback in the form of a green checkmark (correct response) or a red cross (incorrect response or no response) shown for 250 ms. The total trial duration was 8 s. Trials were grouped into blocks of 4 trials of the same condition (32 s total). Each scanning run consisted of 12 blocks (6 per condition) and 4 blocks of a baseline fixation screen (16 s each), for a total run duration of 448 s. Condition order was counterbalanced across runs. Each participant completed 2 runs.

#### Story listening task (C1)

C1 completed several functional MRI tasks. Here, we describe the language task that was analyzed for the current study. The fMRI stimuli consisted of 40 pairs of two-sentence audio-based “stories”. The pairs of stories were identical apart from 1 to 3 words in the second sentence (target), which rendered half of the stories less coherent given the preceding sentence (context) (e.g., “Lindsey loved warm weather. Summer was her favorite season.” vs. “Lindsey loved warm weather. Winter was her favorite season.”). To align as much as possible with EG’s results (which focused on language comprehension, and not the coherence of stories) we only included trials in which the context and target sentences were coherent. The stories were recorded by a native speaker of Standard American English. Stories were presented in an event-related fMRI design. In each trial, the context sentence was presented followed by a jittered inter-stimulus interval and then the target sentence. Trials were separated by a jittered inter-trial interval. In some cases, the target would be followed by a catch trial, in which participants would hear a statement regarding the story that they had just heard (e.g., “The story mentioned a math problem.”: “TRUE or FALSE?”). Participants responded by pressing a button with their dominant hand. The purpose of the catch trials was to keep participants engaged in the task. Participants received a short training session outside the scanner to familiarize them with the task. Inside the scanner, they were instructed to listen carefully to the stories—no explicit semantic judgment task was required. The fMRI task was split into two runs of ∼10 min each.

### fMRI data acquisition

Structural and functional data from all participants scanned at MIT were collected on a whole-body, 3 T, Siemens Trio scanner with a 32-channel head coil, at the Athinoula A. Martinos Imaging Center at the McGovern Institute for Brain Research at MIT. T1-weighted structural images were collected in 176 sagittal slices with 1 mm isotropic voxels (TR = 2530 ms, TE = 3.48 ms). Functional, blood oxygenation level dependent (BOLD), data were acquired using an EPI sequence (with a 90° flip angle and using GRAPPA with an acceleration factor of 2), with the following acquisition parameters: thirty-one 4-mm-thick near-axial slices acquired in interleaved order (with 10% distance factor), 2.1 mm × 2.1 mm in-plane resolution, field of view of in the phase encoding (A > P) direction 200 mm and matrix size 96 × 96, TR = 2000 ms and TE = 30 ms. Prospective acquisition correction (Thesen et al., 2000) was used to adjust the positions of the gradients based on the participant’s motion from the previous TR. The first 10 s of each run were excluded to allow for steady-state magnetization.

Structural and functional data from C1 were collected on a 3 T Siemens Prisma scanner with a 32-channel head coil on the medical campus of Northwestern University. A T1-weighted structural scan was acquired for C1 with TR = 2300 ms, TE = 2.99 ms, flip angle = 7°, inversion time = 900 ms, and 208 contiguous sagittal slices (slice thickness = 0.8 mm, voxel size = 0.8 × 0.8 × 0.8 mm^3^, matrix size = 320 × 320). The functional T2*-weighted images were acquired using an echo-planar sequence with TR = 2000 ms, TE = 25 ms, flip angle = 80°, and 64 axial slices in interleaved order (slice thickness = 2 mm, voxel size = 2 × 2 × 2 mm^3^, matrix size = 104 × 98).

### fMRI data preprocessing

Data were preprocessed with fMRIPrep 23.1.4 (Esteban et al., 2019). Functional images were slice-time corrected, realigned, and co-registered to the T1w reference image. Gray matter (GM), white matter (WM), cerebrospinal fluid (CSF), and whole-brain masks were generated in the anatomical native space. Functional images and masks were then normalized to the MNI152NLin2009cAsym space (resampled to 2 mm). Preprocessed images were smoothed at 6 mm FWHM using 3dmerge from AFNI 20.3.01 (Cox and Hyde, 1997). Individual volumes were marked as outliers if framewise displacement was greater than 0.5 mm.

### fMRI data modeling and statistical analysis

#### First-level modeling

First-level modeling was performed using SPM12. For each participant, task conditions were included as separate regressors in a general linear model (GLM; e.g., for the language task, the conditions included Sentences and Nonwords; for the spatial WM task: Hard and Easy). We created the following contrasts for each participant: language: *Sentences > baseline*, *Nonwords > baseline*, *Sentences > Nonwords* (the critical contrast of interest); spatial WM: *Hard > baseline*, *Easy > baseline*, *Hard > Easy* (the critical contrast of interest).

For C1, task conditions (Coherent Context and Target Sentences) were included as separate regressors in a general linear model. Incoherent context and target sentences, catch trials, responses from catch trials, and inter-stimulus intervals were modeled as regressors of no interest. The contrast of interest was *Coherent Context Sentences + Coherent Target Sentences > baseline (i.e., Spoken Stories > baseline)*. Outlier volumes were included in the GLM as covariates of no interest.

For both cases, the masking threshold was set to -Inf to prevent SPM from excluding voxels in the cerebellum due to lower signal from these regions, and an explicit whole-brain mask was used (Thürling et al., 2012).

#### Group-level analyses

To visualize the average response across participants, group-level analyses were performed using SPM12. First-level maps for the main contrasts of interest (Language: Sentences > Nonwords; WM: Hard > Easy; Spoken Stories > baseline for C1) were brought up to the second level. One-sample t-tests were conducted for each contrast of interest. Age and gender were included as covariates.

#### Lateralization index (LI)

To quantify language lateralization in the cerebellum, we computed a cerebellar lateralization index for each participant as 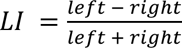, where left and right were the mean Sentences > Nonwords (or Sentences > baseline) contrast values (thresholded to retain positive values; the Spoken Stories > baseline contrast was used for C1) in the left and right hemispheres, respectively. LI ranges from -1 (fully right-lateralized) to 1 (fully left-lateralized). In addition to this magnitude-based approach of computing LI, we also used a voxel-count-based approach to further examine C1’s language lateralization (see **Supplementary Method Section 2**).

#### Spatial correlation

To compare EG’s cerebellar language activation to a typical activation landscape, we used a whole-brain probabilistic language system map derived from 806 TD participants who performed the same or a similar language localizer paradigm as the one used here (Lipkin et al., 2022). All analyses were restricted to cerebellar voxels defined by the SUIT anatomical atlas (Diedrichsen et al., 2009). Within this mask, we extracted contrast values from all voxels with positive, finite values in EG’s map and in each individual TD’s map for the Sentences > Nonwords contrast and in C1’s map for the Spoken Stories > baseline contrast, and then computed a Pearson correlation coefficient (a) between EG’s map and the probabilistic map, (b) between C1’s map and the probabilistic map, and (c) between each TD map and the probabilistic map, to create a baseline distribution. Because EG’s language system is lateralized to the right hemisphere (as originally reported in Tuckute et al., 2022), we compared her activations to the left-right flipped version of the probabilistic map.

#### Identification of subject-specific language-responsive regions

To identify language-responsive regions within the cerebellum in each individual, we defined EG’s and each TD individual’s language-responsive regions using parcels that correspond to typical locations of language activations in a large group of participants. In particular, we used the cerebellar parcels created from cerebellar language activation maps in n = 754 participants, all of whom completed a language localizer task (Casto et al., 2026). This set of parcels includes five parcels in the right cerebellar hemisphere, located in lateral lobule VI (Lang_cereb_1), posterior lobule VI and Crus I (Lang_cereb_2), lobules Crus I, Crus II, and VIIb (Lang_cereb_3), lobules VIIb and VIIIa (Lang_cereb_4), and lobule IX (Lang_cereb_5); and four parcels in the left cerebellar hemisphere, located in lateral lobule VI (Lang_cereb_1), posterior lobule VI (Lang_cereb_2), lobules VI, Crus I, Crus II, and VIIb (Lang_cereb_3), and lobules VIIb and VIIIa (Lang_cereb_4). Note that these parcels were created from a group-constrained subject specific (GCSS) analysis applied exclusively to the cerebellum. A similar analysis applied to the whole brain results in 4 parcels (see Casto et al., 2026). Restricting the analysis to the cerebellum provides greater anatomical resolution, revealing additional subdivisions that are not resolved in the whole-brain analysis. For EG, we used the RH set for her left cerebellar hemisphere, and the LH set for her right cerebellar hemisphere given the flipped hemispheric bias.

To extract language responses for each participant, we identified the 10% of voxels with the strongest Sentences > Nonwords response (based on the contrast values) in one run of the language localizer as language-responsive functional regions of interest (fROIs), and extracted responses to the Sentences > Nonwords contrast from those fROIs using held-out runs. To investigate language selectivity, we additionally extracted responses to the Hard > Easy contrast in the same language fROIs from all runs of the spatial WM task. For example, for EG, who completed four runs each of the language task and the spatial WM task, the Sentences > Nonwords contrast in Run 1 was used to define language-responsive fROIs. Language responses were then extracted from Runs 2-4 of the language task, and WM responses in the same regions were extracted from all four runs of the spatial WM task. This procedure was performed iteratively for each of the remaining three runs.

Language selectivity in each language region was computed by comparing the fMRI response to the Sentences > Nonwords contrast to the fMRI response to the Hard > Easy contrast in the spatial WM task.

Given that both C1’s LH and RH language activation patterns were more similar to a typical right cerebellum (**Figure 5B inset**), we used the RH set of the cerebellar parcels for both of her cerebellar hemispheres. The Spoken Stories > baseline contrast was used to define language-responsive fROIs and extract language responses.

*Statistical analyses.* We used statistical methods designed for single-case studies, following Tuckute et al. (2022). To compare the lateralization index, response magnitudes, and spatial correlations between EG and the TD group, we used the Crawford–Howell modified *t*-test (two-tailed; Crawford and Garthwaite, 2007) as implemented by the crawford.test() function in the *psycho* package in R. This approach evaluates whether a single individual’s value is significantly different from a control sample while appropriately accounting for uncertainty in the control group mean and variance, deriving the point estimate and credible interval via 10,000 Monte Carlo simulations.

## Supporting information

Suppmental Figures and Methods

## Acknowledgements

We would like to thank our participants, EG and C1, for their time and patience. We also acknowledge the contributions of former and current EvLab members for contributions to the fMRI data collection efforts in EG over the years. We would like to thank Steven L. Small for help with C1’s imaging data and for discussions and comments on the analyses and manuscript. We would also like to thank Elissa L. Newport for sharing contrast maps of patients with left-hemispheric perinatal stroke for qualitative examination.

## Author information

### Contributions

B.W.: conceptualization, methodology, formal analysis, visualization, writing (original draft), and writing (review and editing). G.T.: data curation of EG’s data and writing (review and editing). H.K.: data acquisition of EG’s data, data curation of EG’s data, and writing (review and editing). S.S.A.: acquisition of C1’s data, curation of C1’s data, and writing (review and editing). S.G.-M.: acquisition of C1’s data and writing (review and editing). S.C.L.: acquisition of C1’s data and writing (review and editing). E.F.: writing (review and editing). A.M.D.: conceptualization, methodology, validation, supervision, funding acquisition, and writing (review and editing).

