## Supplementary material for "The cerebellum specializes for language even in the absence of contralateral neocortical inputs": Suppmental Figures and Methods

THIS PAPER IS A PREPRINT: IT HAS NOT YET BEEN PEER-REVIEWED.

### Supplementary Information

#### Supplementary Figure 1. Neocortical activation patterns for language in typically-developing participants and EG.

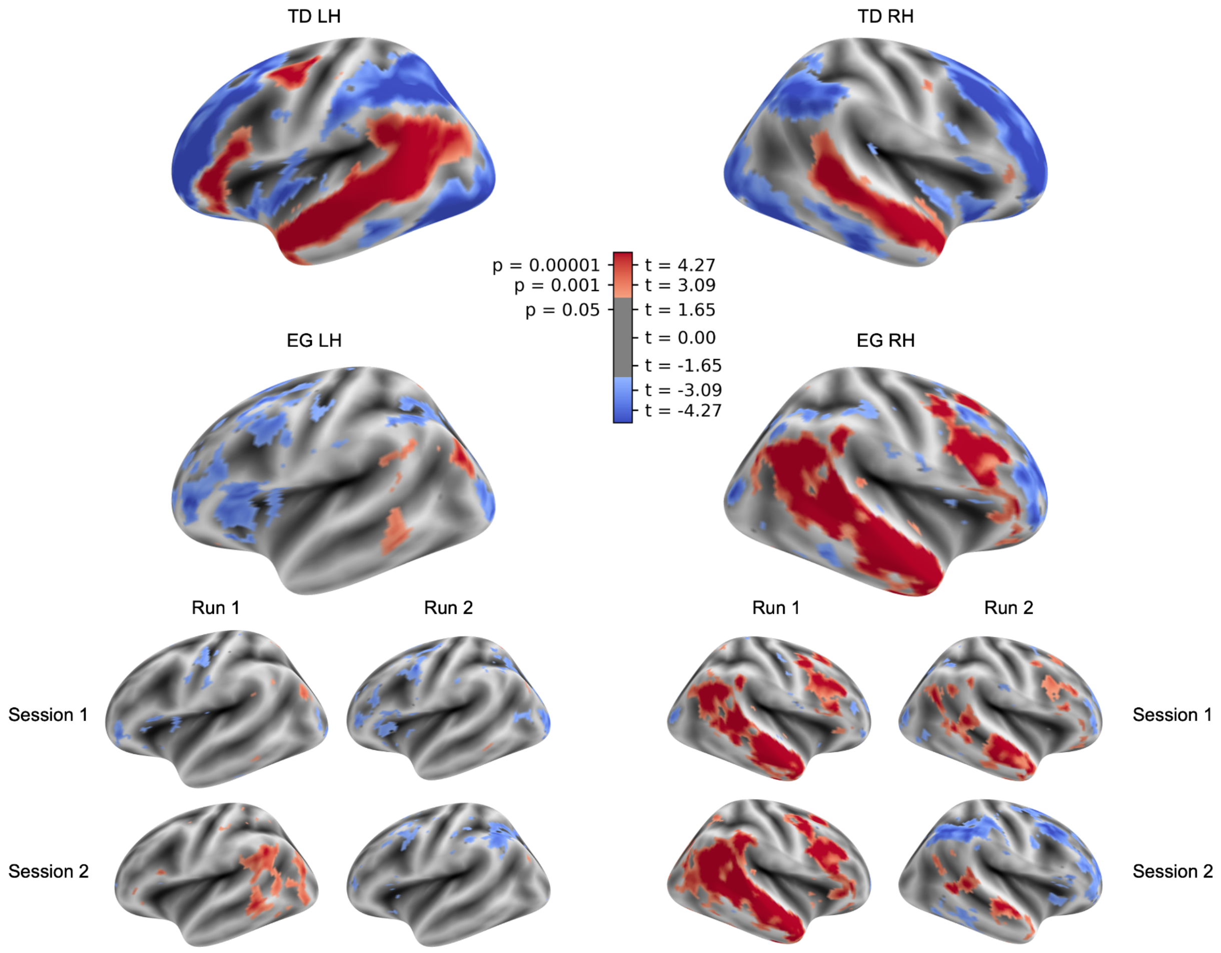


**Supplementary Figure 1. Neocortical activation patterns for language in typically-developing participants and EG. Top.** Group-level t-statistic map for the Sentences > baseline contrast in the language localizer task. TD individuals showed a left-lateralized language network. EG's language network is right-lateralized. Note that across runs, a cluster in the angular gyrus showed a response on this contrast but was not language-responsive (see **Figure S2**). **Bottom.** Responses to Sentences > Nonwords in each of four runs across two sessions. EG showed a strongly right-lateralized neocortical language network. As shown in Tuckute et al. (2022), though anatomically intact, EG's inferior frontal regions ipsilateral to the missing temporal lobe did not show language sensitivity (Sentences > Nonwords).

THIS PAPER IS A PREPRINT: IT HAS NOT YET BEEN PEER-REVIEWED.

**Supplementary Figure 2. Response and connectivity profiles of the left angular gyrus (AG) in EG.**

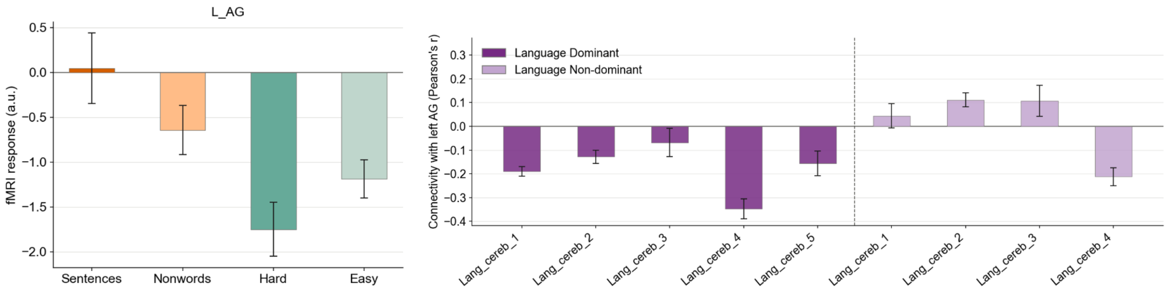


**Supplementary Figure 2. Response and connectivity profiles of the left angular gyrus (AG) in EG. Left**. Response magnitudes for all conditions in the language (Sentences and Nonwords) and spatial working memory tasks (Hard and Easy) within the left angular gyrus (shown in **Figure S1**; cluster extending to the left inferior parietal lobule in some runs). This region did not reliably activate above baseline for Sentences. **Right.** Overall, in EG, resting-state functional connectivity strengths between left AG and cerebellar language regions were negative or low. Remarkably, EG's non-dominant Lang_cereb_4, which showed language selectivity, was not correlated with the left AG.

THIS PAPER IS A PREPRINT: IT HAS NOT YET BEEN PEER-REVIEWED.

**Supplementary Figure 3. Activation patterns for language in C1's neocortex and left cerebellum.**

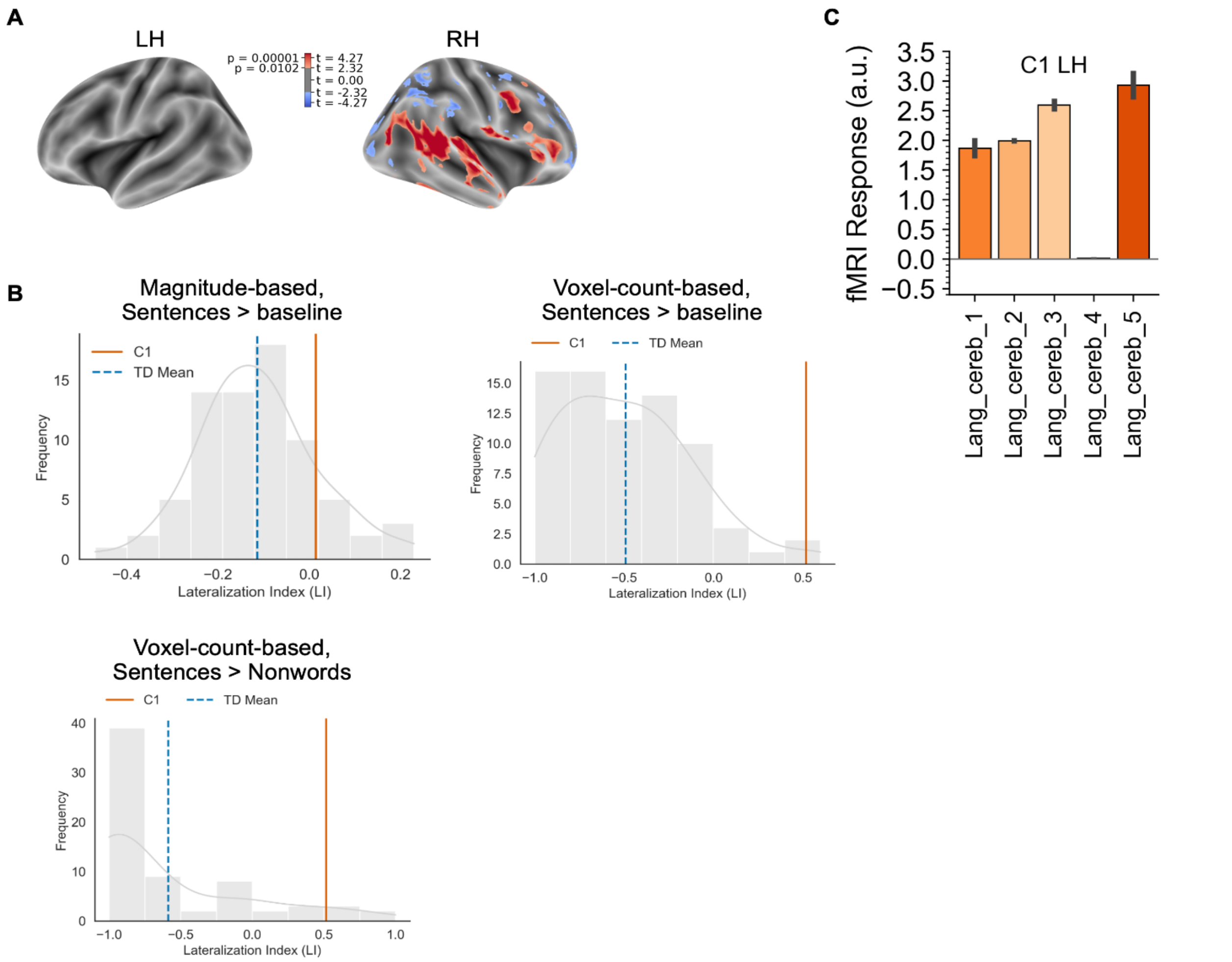


**Supplementary Figure 3. Activation patterns for language in C1's neocortex and left cerebellum. A.** C1's t-statistic map for the Spoken Stories > baseline contrast in the story listening task. **B.** *Top Left.* Distribution of the *magnitude-based* lateralization indices (LIs; see **Methods**) in TDs' activations for the Sentences > baseline contrast. *Top Right.* Distribution of the voxel-count-based LIs (see **Supplementary Method Section 2**) in TDs' activations for the Sentences > baseline contrast. *Bottom Left.* Distribution of the voxel-count-based LIs (see **Supplementary Method Section 2**) in TDs' activations for the Sentences > Nonwords contrast. C1's LI for the Spoken Stories > baseline contrast is shown in orange, and the mean TD LI is shown in blue. **C.** Responses to the Spoken Stories > baseline contrast in C1's left cerebellar regions. Given that C1's LH language activation pattern is more similar to a typical RH pattern (see **Figure 5B inset**), the responses were extracted from TDs' language-dominant (right) cerebellar parcels. The error bars indicate the standard error of the mean across experimental runs.

#### Supplementary Method Section 1. Characterization of the response and connectivity profiles of the left angular gyrus in EG.

Extracting responses from EG's left hemisphere is challenging given the atypical anatomy. Because fROI-based analyses rely on parcels (defined in TDs), and these are difficult to reliably warp to the atypical anatomy of EG's left hemisphere, we used a different data-driven approach to determine whether activation seen here was language-responsive. We first identified reliable clusters across runs, and identified the peak voxel in each cluster — only one cluster emerged reliably in most runs. We then drew a spherical ROI around the peak voxels in this cluster in each run, and extracted language responses from this ROI in held-out runs. We found that this area was not language-responsive in EG, as it did not respond above baseline for Sentences.

*Clusterization*. Each of the Sentences > Nonwords contrast maps in EG was masked to the neocortex and thresholded at $t \geq 2.33$. In each resulting map, clusters were identified using SPM12's spm_bwlabel with 26-connectivity (NN3), and clusters smaller than 10 voxels were discarded.

*ROI definition.* In each contrast map, a 4-mm-radius sphere was centered on the peak voxel of the cluster. Responses from held-out runs were extracted from the spherical ROI.

*Resting-state task.* EG was instructed to close her eyes and let her mind wander for 5 minutes while trying her best to stay awake. The projector was turned off and the lights in the room were dimmed. EG completed a resting-state task in Session 1.

*Resting-state functional connectivity analyses.* Resting-state functional connectivity analyses were conducted using the CONN-fMRI Functional Connectivity toolbox (version 20b) (Whitfield-Gabrieli and Nieto-Castanon, 2012). Functional data were band-pass filtered (0.01–0.1 Hz). Other potential confounds such as white matter, cerebrospinal fluid (CSF), and movement parameters were removed from images. Global signal removal was not applied. ROI-to-ROI connectivity was computed as the bivariate Pearson correlation between each ROI pair (left AG fROI defined in four runs × nine cerebellar language fROIs defined in four runs).

#### Supplementary Method Section 2. Description of the voxel-count-based lateralization index.

*Lateralization index (LI).* To quantify language lateralization in the cerebellum, we extracted the number of significant voxels ($p < 0.001$ uncorrected) from the whole cerebellum and computed a cerebellar lateralization index for each participant as $LI = \frac{left - right}{left + right}$, where left and right were the number of significant voxels in the Sentences > Nonwords (or Sentences > baseline) *t*-statistic map (the Spoken Stories > baseline *t*-statistic map was used for C1) in the left and right hemispheres, respectively. LI ranges from -1 (fully right-lateralized) to 1 (fully left-lateralized).

THIS PAPER IS A PREPRINT: IT HAS NOT YET BEEN PEER-REVIEWED.
